# Limits of aerobic metabolism in cancer cells

**DOI:** 10.1101/020461

**Authors:** Alexei Vazquez

## Abstract

Cancer cells exhibit high rates of aerobic glycolysis and glutaminolysis. Aerobic glycolysis can provide energy and glutaminolysis can provide carbon for anaplerosis and reductive carboxylation to citrate. However, all these metabolic requirements could be in principle satisfied from glucose. Energy can be generated from oxidative phosphorylation (OxPhos) of glucose, anaplerosis can be accomplished using pyruvate carboxylate and citrate can be derived from glucose. Here we investigate why cancer cells do not satisfy their metabolic demands using aerobic biosynthesis from glucose. Based on the typical composition of a mammalian cell we quantify the energy demand and the OxPhos burden of cell biosynthesis from glucose. Our calculation demonstrates that aerobic growth from glucose is feasible up to a minimum doubling time that is proportional to the OxPhos burden and inversely proportional to the mitochondria OxPhos capacity. To grow faster cancer cells must activate aerobic glycolysis for energy generation and uncouple NADH generation from biosynthesis. To uncouple biosynthesis from NADH generation cancer cells can synthesize lipids from carbon sources that do not produce NADH in their catabolism, including acetate and the amino acids glutamate, glutamine, phenylalanine and tyrosine. Finally, we show that cancer cell lines commonly used in cancer research have an OxPhos capacity that is insufficient to support aerobic biosynthesis from glucose. We conclude that selection for high rate of biosynthesis implies a selection for aerobic glycolysis and uncoupling biosynthesis from NADH generation. Any defect or perturbation reducing the OxPhos capacity will exacerbate this selection.

## Introduction

Cancer cells exhibit high rates of glucose fermentation to lactate even when growing in aerobic conditions, a phenotype known as aerobic glycolysis or the Warburg effect. The Warburg effect is accompanied by other metabolic alterations, particularly increased glutamine utilization (1-4) and reductive carboxylation of glutamine to the lipid precursor AcCoA (5). More recently, acetate has been shown to be another important source of AcCoA in cancer cells (6-9) and its contribution increases under oxygen limitation (6).

Given that fermentation of glucose is the default pathway under anaerobic growth, Warburg originally thought that aerobic glycolysis should be rooted on mitochondrial defects (10, 11). However, cells from healthy mammalian tissues also manifest aerobic glycolysis and glutaminolysis during fast growth (12-15). Furthermore, aerobic glycolysis is observed with concomitant high rates of respiratory metabolism in cancer (15) and muscle (16) cells.

Quantitative models of cell metabolism have been used to investigate the cause of aerobic glycolysis and related metabolic phenotypes. The postulation of a maximum oxygen consumption rate results in model predictions mimicking aerobic glycolysis in yeast (17) and the Warburg effect (unpublished data). However, under aerobic conditions, further increase of the oxygenation level does not abrogate aerobic glycolysis (16, 18). Simulations of genome scale models of mammalian cell metabolism also predict the utilization of glutamine at high proliferation rates (19, 20). However, we do not understand what aspects of the model are responsible for that prediction.

## Results

### Aerobic growth from glucose

To gain a better understanding of cell metabolism as a function of the growth metabolic demand we performed a back-of-the-envelope calculation focusing on the major biomass components of mammalian cells. The dry weight of mammalian cells is mostly composed of protein, lipids and polysaccharides (Table 1, Table S1). Based on this composition we can estimate the biosynthetic demand for precursor metabolites and the energy that is necessary to duplicate the cell content. The duplication of a typical mammalian cell requires about 1.1 mol/L of amino acids for protein synthesis, 1.2 mol/L of AcCoA for lipid synthesis and an energy demand of *b*_demand_=6.0 mol/L of ATP (Table 1).

**Table 1:**
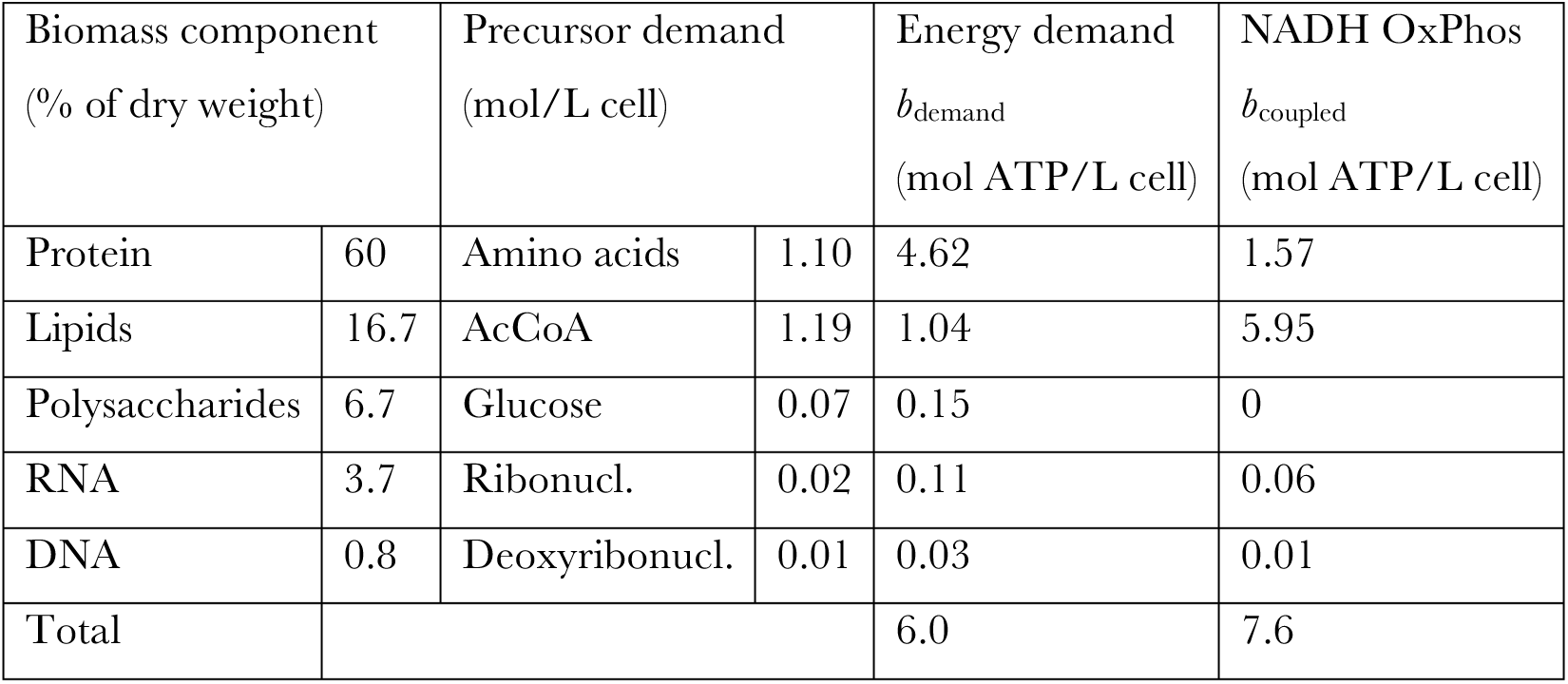
Composition and growth requirements of a typical mammalian cell. The NADH OxPhos column reports the energy that can be generated coupled to OxPhos of the NADH produced by biosynthetic pathways from glucose.

The synthesis of biomass precursors from glucose makes use of different dehydrogenases producing/consuming NADH (Fig. 1). The synthesis of some amino acids involves transamination coupled with glutamate to α-ketoglutarate conversion. α-ketoglutarate can be converted back to glutamate via reverse glutamate dehydrogenase consuming NADH. When all these contributions are taken into account we obtain a net production of NADH coupled to the biosynthesis of precursor metabolites from glucose. The NADH produced can be oxidized in the mitochondria coupled to ATP generation. Assuming a 2.5 ATP/NADH yield we obtain that mammalian cells can produce *b*_coupled_=7.6 mol ATP/L coupled to NADH production by biosynthetic pathways (NADH OxPhos, Table 1).

**Figure 1:**
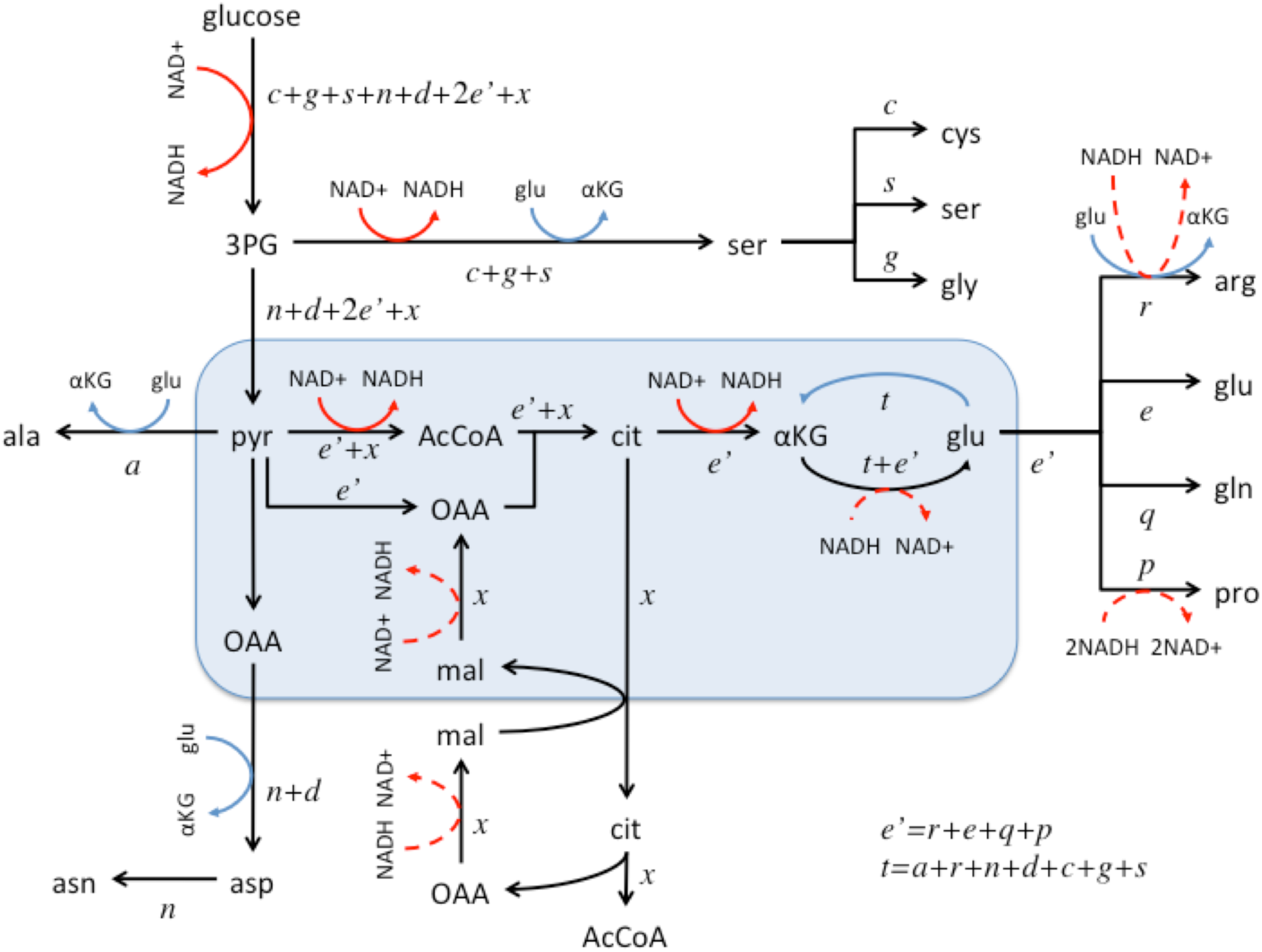
Synthesis of precursor metabolites from glucose. The text in italics represents the precursor requirements to duplicate a cell, using the one letter nomenclature for amino acids and *x* for AcCoA. Dehydrogenase steps are highlighted with red arrows (solid forward and dashed reverse) and transaminase steps with blue arrows. ATP and other cofactors have been omitted for simplicity.

The aerobic biosynthesis of a cell from glucose relies on the capacity of mitochondria to generate the energy required for biosynthesis and to oxidize the NADH generated through biosynthesis. In the following we calculate how fast a cell can proliferate given a specified mitochondrial content. If *a* is the growth-independent energy demand for cell maintenance, *b* is the OxPhos demand to duplicate the cell content, *T* is the cell doubling time, *r*_M_ is the OxPhos capacity of mitochondria (mol ATP/L of mitochondria) and *ϕ*_M_ is the mitochondria volume fraction (L mitochondria/L cell) then the following constraint must be satisfied

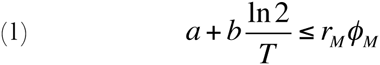

If cells proliferate too fast, shortening the doubling time (left hand side of equation 1), they will overcome the maximum OxPhos capacity (right hand side of equation 1). From equation 1 we can determine the minimum doubling time that is supported by aerobic biosynthesis from glucose

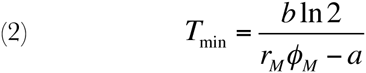

The mitochondria OxPhos capacity (*r*_M_) has been measured for isolated mitochondria (Fig. 2, Table S2). It can be as high as 0.38 mol ATP/L mitochondria/min in muscle cells. The mitochondria of yeast cells is as efficient, with *r*_M_=0.29 mol ATP/L mitochondria/min. However, the mitochondria of cancer cells is 10 fold less efficient, with *r*_M_=0.042-0.068 mol ATP/L mitochondria/min.

**Figure 2:**
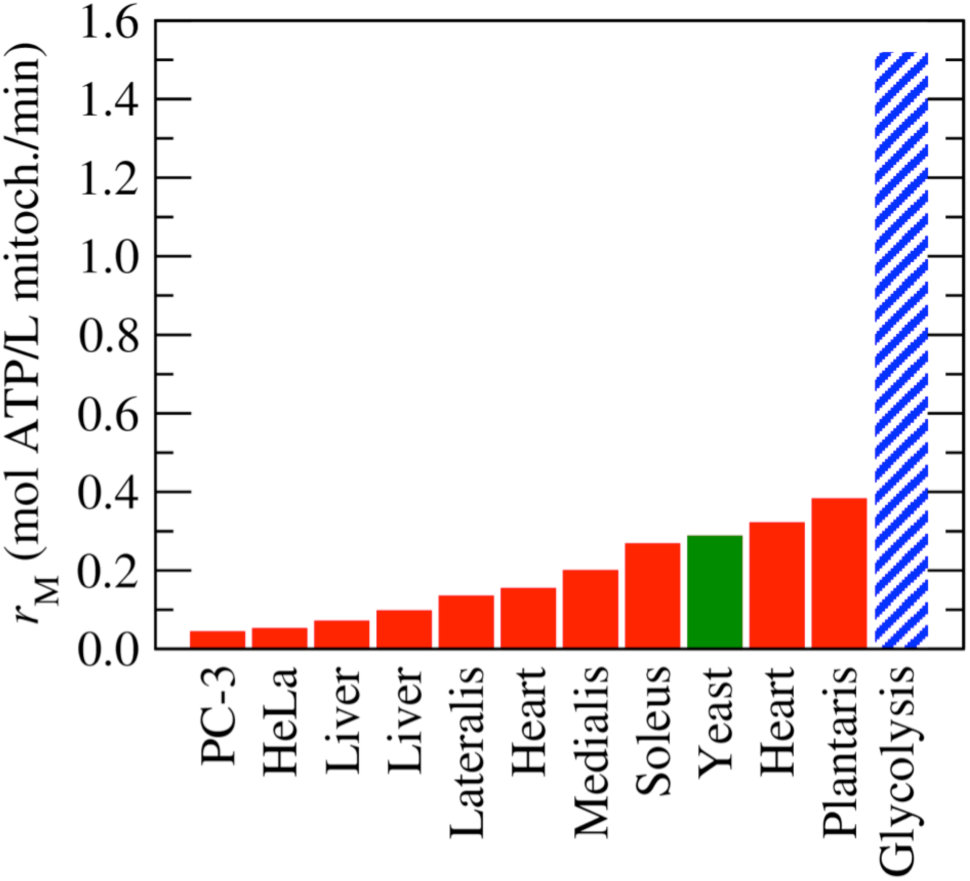
Mitochondria capacity. Mitochondria intrinsic capacity (per volume of mitochondria) for ATP generation, based on data reported in the literature (Table S2). Red represents data for different mammalian tissues and cell lines. Green represents data for yeast. As a reference, in dashed-blue we highlight the intrinsic capacity of glycolysis.

### Growth phases

The growth phase diagram implied by equation (2) is shown in Figure 3. The top right corner corresponds with to a growth phase where cells can proliferate producing all the energy from OxPhos and can synthesize precursor metabolites from glucose (facultative OxPhos and facultative coupling of biosynthesis to NADH production, FF phase). We notice that “they can” does not imply “they must”. A cell line may have a doubling time and mitochondria content in that region, but it may have up-regulated aerobic glycolysis due to genetic alterations. Cells may also utilize alternative carbon sources due to overexpression of transport systems, reducing the NADH production coupled to biosynthesis. In such cases we can truly speak of an inefficient/wasteful/accidental phenotype. The other extreme is the grey zone highlighted in Fig. 3, where the energy demand associated with cell maintenance exceeds the OxPhos capacity (*a*>*r*_*MϕM*_ in equation 2). In this region cells cannot survive without activating aerobic glycolysis.

**Figure 3:**
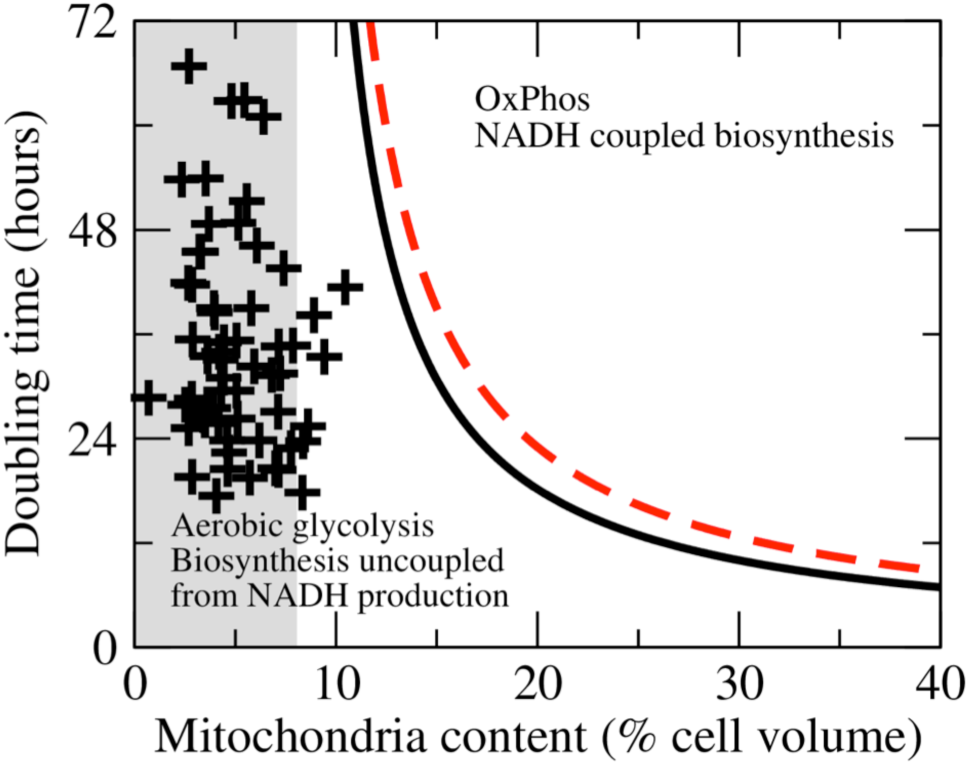
Growth phase diagram. phases as a function of cell mitochondrial content. The Growth doubling time and black-solid obtained using equation 2, setting the demand equal to the energy demand duplication (*b*=*b*_demand_). The red dashed line is OxPhos of cell line is obtained using equation 2, setting the OxPhos demand equal to the NADH OxPhos of cell duplication (*b*=*b*_coupled_). The grey area demarks the region where the OxPhos capacity is insufficient to satisfy the energy demand for cell maintenance. The plusses represent 60 cancer cell lines from the NCI60 panel.

In mammalian cells the energy that can be generated from OxPhos of the NADH produced in biosynthetic pathways exceeds the energy demand of biosynthesis (*b*_demand_<*b*_coupled_, Table 1). As cells decrease their doubling time and/or decrease their mitochondrial content, they will first be limited by the inability of mitochondria to turnover the NADH produced coupled to the biosynthesis of precursor metabolites (Fig. 3, red-dashed line). To proliferate faster, shortening the doubling time, mammalian cells need to uncouple NADH production from the synthesis of precursor metabolites (facultative OxPhos and obligatory uncoupling of biosynthesis from NADH production, FO phase). In the FO phase mammalian cells can satisfy their energy demand from OxPhos, until the energy demand of biosynthesis reaches the OxPhos capacity (Fig. 3, black-solid line). Below the black-solid line mammalian cells cannot satisfy their energy demand using OxPhos and they must switch on aerobic glycolysis (obligatory aerobic glycolysis and obligatory uncoupling of biosynthesis from NADH production, OO phase). In this case aerobic glycolysis is not an inefficient/wasteful/accidental phenotype, it is the only choice left to proliferate faster.

To determine the location of cancer cells in the growth phase diagram of Fig. 3, we focus on the NCI60 panel of tumour derived cell lines. The doubling times for these cell lines has been reported by the NCI Developmental Therapeutics Program and we have estimated their mitochondrial content. Each cross in Fig. 3 represents a cell line in the NCI60 panel. Most cell lines fall in the grey area, where OxPhos cannot supply the energy requirements of cell maintenance and aerobic glycolysis must be activated. Indeed, these cell lines convert approximately 70% of the imported glucose to lactate (21, 22). For these cancer cells we can conclude that aerobic glycolysis is a consequence of a limited OxPhos capacity. Metabolic flux estimations indicate that these cells generate ATP from OxPhos in amounts comparable to aerobic glycolysis (22). The observation of this mixed aerobic-glycolysis/OxPhos phenotype does not contradict our conclusion that aerobic glycolysis is obligatory for these cells. Simply the energy generated from OxPhos falls short of the energy demand.

### Uncoupling NADH production from biosynthesis

Biosynthesis can be uncoupled from NADH generation in different ways. NADH can be turnover (NADH →NAD+) coupled to NADPH production (NADP+→NADPH), using combinations of cytosolic/mitochondrial NAD+ and NADP+ dehydrogenases and the mitochondria transhydrogenase. We notice that the conversion of NADH to NADPH by the transhydrogenase transports protons from the mitochondria matrix to the cytosol, uncoupling OxPhos from ATP generation. Therefore, the NADH turnover by the transhydrogenase comes at expenses of exacerbating the limited OxPhos for energy production

As an alternative, cells can switch to other carbon sources to reduce NADH production. The limited OxPhos capacity dictates which carbon sources are more suitable to uncouple NADH production from biosynthesis. Certainly for each amino acid the amino acid itself is a suitable alternative carbon source. The choice is in principle more redundant for AcCoA (Fig. 4a). Acetate can be used with the extra cost of 1 ATP for the acetyl CoA ligase step. On the other hand, the β-oxidation of fatty acids generates NADH and FADH_2_ and therefore it does not reduce the NADH generation burden. AcCoA could also be produced from amino acids. However, in most cases NADH is generated (Fig. 4a, grey lines). The catabolism of isoleucine, leucine, lysine, methionine, tryptophan, and valine requires one or more dehydrogenase steps generating NADH. Alanine, cysteine, glycine, threonine and serine can be converted to pyruvate, which still requires pyruvate dehydrogenase to produce AcCoa (Fig. 1). Aspartate and asparagine can be converted to oxaloacetate, which still requires a source of AcCoA for citrate synthesis or conversion to pyruvate, again requiring pyruvate dehydrogenase to produce AcCoa. Finally, arginine and proline can be converted to glutamate but NADH is generated in the process.

**Figure 4:**
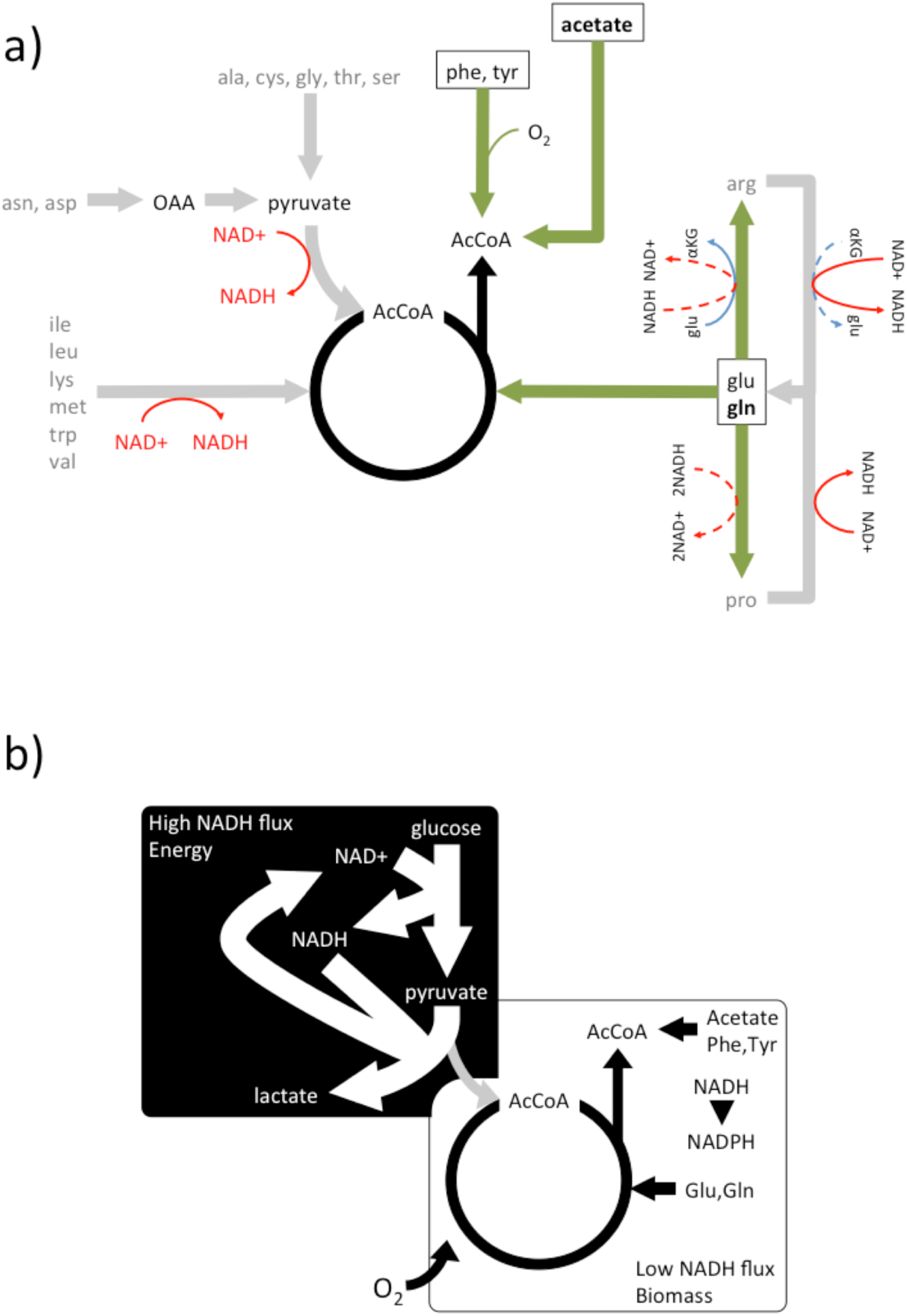
Metabolism under a limited OxPhos capacity. a) Synthesis of AcCoA from alternative carbon sources. In grey we highlight carbon sources that result in NADH production. The boxes highlight the carbon sources that uncouple AcCoA synthesis from NADH production. b) Compartmentalization of eukaryote aerobic cell metabolism in the context of a limited OxPhos capacity.

There are only two groups of amino acids that can be converted to citrate without NADH production. The first group includes glutamate and glutamine. Glutamate can be converted to citrate via reductive carboxylation. In this pathway the NAD(P)H production by glutamate dehydrogenase is compensated by the reverse activity of the NAD(P) isocitrate dehydrogenase (Fig. 1). Glutamate can be taken from the medium or generated from glutamine by glutaminase. Interestingly, arginine and proline can be produced from glutamate with concomitant consumption of NADH (Fig. 4a). This could provide an additional mechanism for NADH turnover. The second group is composed of the amino acids phenylalanine and tyrosine, which are converted to acetoacetate and fumarate (Fig. 4a). Acetoacetate yields 2 AcCoa *via* the combined activity of acetoacetate CoA ligase (consuming one ATP) and acetyl-CoA transferase. Fumarate on the other hand is better excreted to avoid additional OxPhos burden. We notice that the catabolism of phenylalanine and tyrosine to acetoacetate consumes 3 and 2 oxygen molecules respectively. Therefore, these amino acids are not suitable to uncouple NADH production from biosynthesis under hypoxia.

### NADPH production

For the sake of simplicity we have omitted the balance of some key co-factors that are also required for cell proliferation. NADPH is a free energy source required in the biosynthesis of some amino acids and of lipids. The pentose phosphate pathway (PPP) is a major source of NADPH production. The PPP is interconnected with the upper part of glycolysis, above the glyceraldehyde 3-phosphate dehydrogenase step. Therefore, the PPP does not change the pyruvate/NADH yield ratio of glycolysis, which is the only feature of glycolysis used to calculate the biosynthetic requirements reported in Table 1. The increase in the NADPH yield just reduces the percentage of glucose carbons incorporated into biomass (1 CO_2_ molecule is released per every 2 NADPH).

NADPH can also be produced *via* one-carbon metabolism using serine as the one-carbon source (23, 24). The synthesis of serine from glucose generates NADH (Fig. 1). Therefore the glucose→serine→one-carbon pathway of NADPH generation is only suitable when cells have not reached their maximum OxPhos capacity. When cells have overcome their OxPhos capacity they could import serine from the media to uncouple NADH production from NADPH production via one-carbon metabolism. To donate the one-carbon unit serine is converted to glycine and glycine could also donate a one-carbon unit *via* the glycine cleavage system. The glycine cleavage system generates NADH and therefore this pathway is only suitable when cells have not reached their maximum OxPhos capacity. Otherwise, glycine is better excreted to avoid additional NADH production.

## Conclusions

When the metabolic demand exceeds the limited OxPhos capacity mammalian cells split their metabolism into two major components (Fig. 4b). There is a component of high NADH flux from aerobic glycolysis (Fig. 4b, black background box). The function of this component is to satisfy the energy demand beyond the OxPhos capacity. There is another component of low NADH flux where precursor metabolites are produced from glucose and other carbon sources (Fig 4b, white background box). The function of this component is to generate precursor metabolites with the minimal NADH generation that can be absorbed by the limited OxpHos capacity. The choice of alternative carbon sources is restricted to those that do not generate NADH in their conversion to precursor metabolites. For generation of AcCoA, this constraint leave cells with the choice of acetate and the amino acids glutamate, glutamine, phenylalanine and tyrosine.

In the context of cancer metabolism it has been often claimed that aerobic glycolysis is a consequence of an increased demand of carbon atoms for biosynthesis of precursor metabolites (25, 26). A more precise statement is that the increased demand for carbon atoms results in an increased demand for NADH turnover when glucose is the carbon source. However, the excretion of lactate does not solve the problem of NADH production coupled to biosynthesis. While it is true that lactate dehydrogenase converts NADH back to NAD+, the carbon atoms associated with this conversion are excreted as lactate and they do not contribute to the formation of biomass precursors. We conclude, as originally postulated by Warburg and supported by previous calculations (27), that the role of aerobic glycolysis is to satisfy the energy demand in the context of limited mitochondria OxPhos capacity.

These conclusions will hold up to small variations in the parameter values required for the analysis. Imprecisions in the demand of energy and NADH turnover may cause a change in the order of the continuous and dashed lines in Figure 3, but the two phases at the top-right and bottom-left corners will remain. The intrinsic mitochondria efficiency of cancer cells may be larger than the values obtained for PC-3 and HeLa cells, causing some cell lines to fall in the growth phase where aerobic glycolysis is not obligatory. Therefore, our conclusion of aerobic glycolysis being obligatory may not necessarily hold for all cancer cells. In spite of these potential issues, this analysis provides a conceptual framework that brings together apparently disconnected metabolic phenotypes of cancer cells.

## Methods

*Cell composition:* The typical biomass composition of mammalian cells was obtained from Ref. (28). The typical composition in terms of precursor metabolites (mol of precursor/L cell) was calculated using the equation

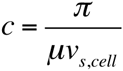

where *π* is the dry weight fraction of the biomass component, *μ* is the molecular weight of the precursor metabolite and *v*_*s,cell*_ is the cell specific volume (L cell/g dry weight). Amino acids are the precursor metabolites of proteins. The average molecular weight of an amino acid in expressed proteins is *μ*_aa_=109 g/mol in mammalian cells (29). The acetyl group of AcCoA is the precursor of lipids. Its average molecular weight in lipids was estimated as two times the molecular weight of CH_2_ (*μ*_ac_=28 g/mol). Ribonucleotides are the precursor metabolites of RNA. The average molecular weight of a ribonucleotide in expressed RNAs is *μ*_rn_=331 g/mol in mammalian cells (29). Deoxyribonucleotides are the precursor metabolites of DNA. The average molecular weight of a deoxyribonucleotide in expressed DNA is *μ*_dn_=332 g/mol in mammalian cells (29). Finally, the cell specific volume is 5 mL/gDW for mammalian cells (30).

*Energy requirements of biosynthesis:* The mammalian cell energy requirement for synthesis of each biomass component from glucose was estimated as the number of ATP molecules required to synthesize the precursor from glucose, based on reported biosynthesis pathways (31), plus the energy required for polymerization (29), times the precursor concentration (Additional File 1, Table S1).

*NADH OxPhos:* The number of NADH molecules produced coupled to the synthesis of each precursor metabolite from glucose was calculated from reported biosynthesis pathways (31). This value was multiplied by a yield of 2.5 ATP/NADH times the precursor concentration reported in Table 1 (see also Table S1).

*Mitochondria OxPhos capacity:* The mitochondria OxPhos capacity *r*_*M*_ was obtained from literature reports (Additional File 1, Table S2).

*Maintenance energy:* The cell maintenance energy demand of mammalian cells was estimated from the data reported for the *LS* mouse cell line (32). *LS* mouse cells require 17 pmol ATP/cell/day for cell maintenance. Dividing by the *LS* cell volume (3.4 pL/cell) we obtain the cell maintenance energy demand per cell volume *a*=3.5 mmol ATP/L cell/min.

*NCI60:* The doubling times for each cell line are reported by the NCI Developmental Therapeutics Program http://dtp.nci.nih.gov/docs/misc/common_files/cell_list.html. Their mitochondrial content was estimated as

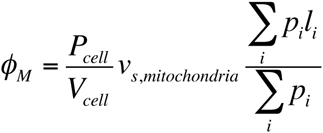

where *P*_*cell*_ is the protein content (g/cell), *V*_*cell*_ is the cell volume (L/cell), *v*_*s,mitochondria*_ is the mitochondrial specific volume (L/g mitochondrial protein), the sums run over all expressed proteins, *p*_*i*_ is the abundance of each protein and *l*_*i*_ =1 if the protein localizes to the mitochondria and 0 otherwise. *P*_*cell*_ and *V*_*cell*_ are reported in Ref. (22). The mitochondria specific volume is *v*_*s,mitochondria*_ = 2.6 mL/g mitochondrial protein (33). The relative protein abundances where obtained from reported proteomic data (34). The protein localization we obtained from the NCBI annotations. *l*_*i*_ was set to 1 for every protein that is annotated to localize to the mitochondria. The estimated mitochondria content is reported in the Additional File 1, Table S3.

## Supplementary material

**Table S1:** Biosynthetic demands of cell biomass duplication from glucose.

| Component | % of cell weight | % of cell dry weight |  |  |  | Energy demand (mol ATP/L cell) |  |  |
| --- | --- | --- | --- | --- | --- | --- | --- | --- |
| Water | 70 |  | Unit | Unit MW (g/mol) | Unit concentration (M) | Per precursor | From glucose | NADH OxPhos (mol ATP/L cell) |
| Ref. | [1] |  |  | [2] |  | [2] |  |  |
| Protein | 18 | 60.0 | Amino acids | 109 | 1.10 | 4.30 | 4.62 | 1.57 |
| Lipids | 5 | 16.7 | Ac | 28 | 1.19 | 0.88 | 1.04 | 5.95 |
| Polysaccharides | 2 | 6.7 | Glucose | 180 | 0.07 | 0.00 | 0.15 | 0.00 |
| RNA | 1.1 | 3.7 | Ribonucle | 331 | 0.02 | 0.40 | 0.06 | 0.06 |
| DNA | 0.25 | 0.8 | Deoxyribonucle | 332 | 0.01 | 1.37 | 0.02 | 0.01 |
| Total | 96.4 | 87.8 |  |  |  |  | 6.0 | 7.6 |
| Synthesis from glucose: (+) produced, (-) consumed |  |  |  |  |  |  |  |  |
| Molecule | Frequency in precursor | ATP/molecule | ATP/precursor | NADH | glu -> akg | Net NADH produced per precursor unit |  |  |
| Ref. | [2] | [3] |  | [3] | [3] |  |  |  |
| ala | 0.087 | 1 | 0.087 | 1 | 1 | 0 |  |  |
| arg | 0.055 | 0 | 0 | 2 | 1 | 0.055 |  |  |
| asn | 0.042 | -1 | -0.042 | 1 | 1 | 0 |  |  |
| asp | 0.052 | 0 | 0 | 1 | 1 | 0 |  |  |
| cys | 0.021 | 0 | 0 | 2 | 1 | 0.021 |  |  |
| glu | 0.056 | 1 | 0.056 | 3 | 0 | 0.168 |  |  |
| gln | 0.047 | 0 | 0 | 3 | 0 | 0.141 |  |  |
| gly | 0.078 | 0 | 0 | 2 | 1 | 0.078 |  |  |
| pro | 0.045 | 0 | 0 | 1 | 0 | 0.045 |  |  |
| ser | 0.062 | 0 | 0 | 2 | 1 | 0.062 |  |  |
| amino acids | 0.545 |  | 0.101 |  |  | 0.570 |  |  |
| AcCoA | 1.00 | 0 | 0 | 2 | 0 | 2 |  |  |
| polysaccharides | 1.00 | -2 | -2 | 0 | 0 | 0 |  |  |
| purine | 0.50 | -6 | -3 | 1 | 0 | 1 |  |  |
| pyrimidine | 0.50 | -3 | -1.5 | 1 | 0 | 1 |  |  |
| RNA |  |  | -2.25 |  |  | 1 |  |  |
| DNA |  |  | -2.25 |  |  | 1 |  |  |
| Parameter | Value | Units | Ref. |  |  |  |  |  |
| Maintenance energy | 17 | pmol/cell/day | [4] |  |  |  |  |  |
| Cell volume | 3.4 | pL | [4] |  |  |  |  |  |
| Maintenance energy | 5 | M/h |  |  |  |  |  |  |
| Cell specific volume | 0.005 | L/g | [5] |  |  |  |  |  |

**Table S2:**
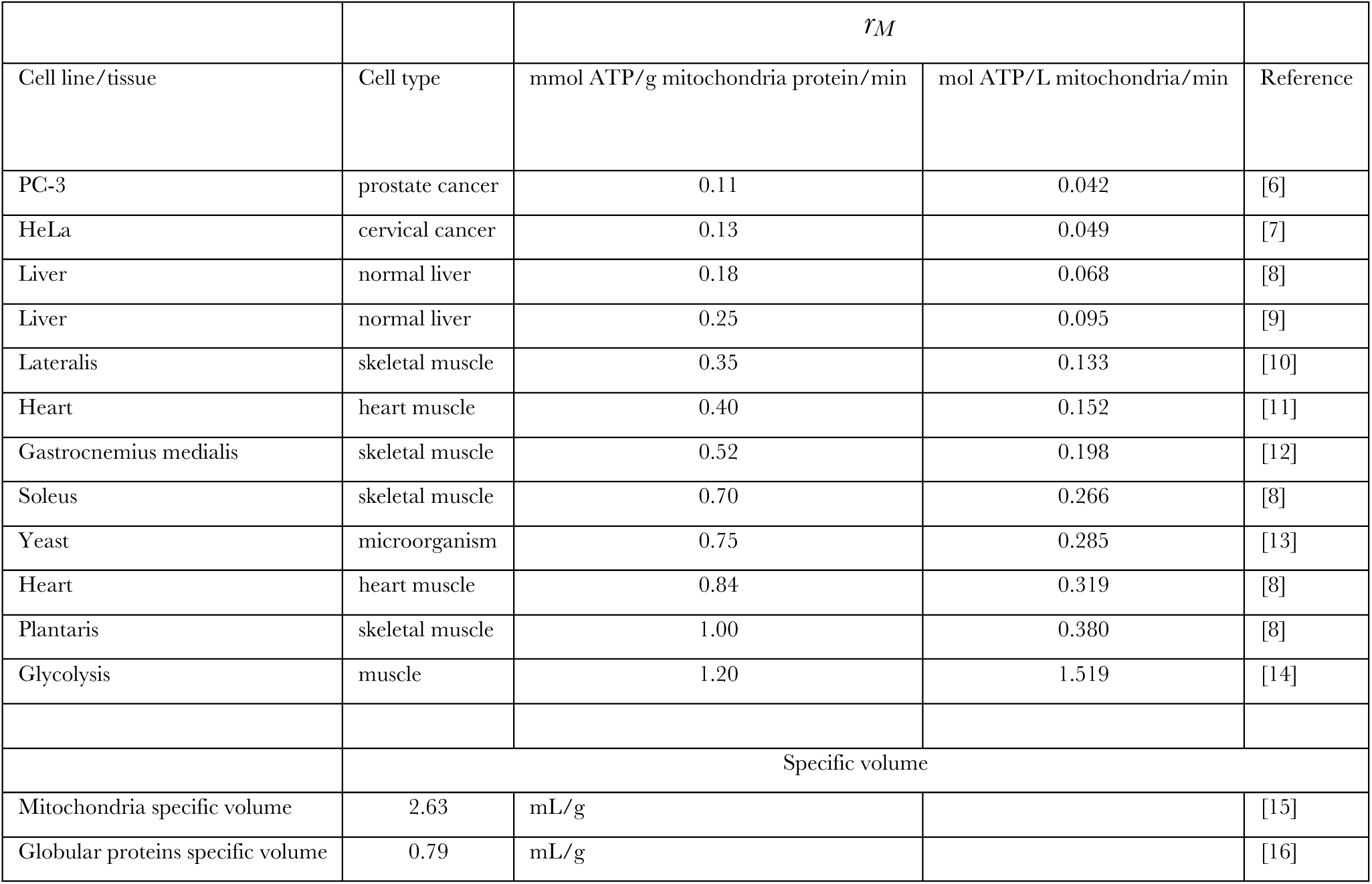
ATP generating capacity of mitochondria from different tissues and cells lines and from glycolysis.

| Cell line/tissue | Cell type | $\dot{r}_M$ | | Reference |
| --- | --- | --- | --- | --- |
|  |  | mmol ATP/g mitochondria protein/min | mol ATP/L mitochondria/min |  |
| PC-3 | prostate cancer | 0.11 | 0.042 | [6] |
| HeLa | cervical cancer | 0.13 | 0.049 | [7] |
| Liver | normal liver | 0.18 | 0.068 | [8] |
| Liver | normal liver | 0.25 | 0.095 | [9] |
| Lateralis | skeletal muscle | 0.35 | 0.133 | [10] |
| Heart | heart muscle | 0.40 | 0.152 | [11] |
| Gastrocnemius medialis | skeletal muscle | 0.52 | 0.198 | [12] |
| Soleus | skeletal muscle | 0.70 | 0.266 | [8] |
| Yeast | microorganism | 0.75 | 0.285 | [13] |
| Heart | heart muscle | 0.84 | 0.319 | [8] |
| Plantaris | skeletal muscle | 1.00 | 0.380 | [8] |
| Glycolysis | muscle | 1.20 | 1.519 | [14] |
|  | Specific volume |  |  |  |
| Mitochondria specific volume | 2.63 | mL/g |  | [15] |
| Globular proteins specific volume | 0.79 | mL/g |  | [16] |

**Table S3:** Estimated protein content of the NCI60 cell lines.

| Cell line | Doubling time | Mitochondrial volume % |
| --- | --- | --- |
| LE:SR | 28.7 | 0.71 |
| BR:HS578T | 53.8 | 2.35 |
| LE:MOLT-4 | 27.9 | 2.52 |
| BR:MDA-MB-231 | 41.9 | 2.67 |
| ME:SK-MEL-5 | 25.2 | 2.68 |
| RE:A498 | 66.8 | 2.71 |
| LE:HL-60 | 28.6 | 2.84 |
| LE:K-562 | 19.6 | 2.86 |
| RE:UO-31 | 41.7 | 2.87 |
| CNS:Sf-539 | 35.4 | 2.89 |
| BR:T-47D | 45.5 | 3.23 |
| ME:SK-MEL-2 | 45.5 | 3.30 |
| LE:CCRF-CEM | 26.7 | 3.41 |
| OV:OVCAR-8 | 26.1 | 3.49 |
| BR:BT-549 | 53.9 | 3.53 |
| LE:RPMI-8226 | 33.5 | 3.63 |
| OV:SK-OV-3 | 48.7 | 3.69 |
| RE:ACHN | 27.5 | 3.89 |
| RE:CAKI-1 | 39 | 3.94 |
| ME:UACC-257 | 38.5 | 3.98 |
| CO:HCT-116 | 17.4 | 4.06 |
| ME:MDA-MB-435 | 25.8 | 4.17 |
| OV:OVCAR-8/ADR | 34 | 4.20 |
| CNS:Sf-268 | 33.1 | 4.29 |
| CNS:Sf-295 | 29.5 | 4.31 |
| OV:IGROV1 | 31 | 4.42 |
| ME:SK-MEL-28 | 35.1 | 4.43 |
| CO:COLO205 | 23.8 | 4.60 |
| ME:LOXIMV1 | 20.5 | 4.62 |
| RE:786-0 | 22.4 | 4.69 |
| CNS:SNB-75 | 62.8 | 4.81 |
| LC:NCI-H322M | 35.3 | 5.07 |
| RE:SN12C | 29.5 | 5.07 |
| ME:M14 | 26.3 | 5.12 |
| OV:OVCAR-5 | 48.8 | 5.15 |
| RE:RXF-393 | 62.9 | 5.46 |
| RE:TK-10 | 51.3 | 5.57 |
| CO:HT29 | 19.5 | 5.72 |
| LC:HOP-62 | 39 | 5.79 |
| PR:DU-145 | 32.3 | 5.94 |
| ME:MALME-3M | 46.2 | 6.07 |
| CNS:U251 | 23.8 | 6.20 |
| LC:NCI-H226 | 61 | 6.41 |
| ME:UACC-62 | 31.3 | 6.82 |
| CO:HCT-15 | 20.6 | 6.99 |
| PR:PC-3 | 27.1 | 7.13 |
| CO:SW-620 | 20.4 | 7.14 |
| CNS:SNB-19 | 34.6 | 7.16 |
| CO:HCC-2998 | 31.5 | 7.23 |
| LC:EKVX | 43.6 | 7.41 |
| LC:A549/ATCC | 22.9 | 7.77 |
| OV:OVCAR-3 | 34.7 | 7.86 |
| LC:HOP-92 | 79.5 | 7.92 |
| LC:NCI-H460 | 17.8 | 8.33 |
| CO:KM12 | 23.7 | 8.36 |
| BR:MCF7 | 25.4 | 8.62 |
| LC:NCI-H522 | 38.2 | 8.91 |
| LC:NCI-H23 | 33.4 | 9.42 |
| OV:OVCAR-4 | 41.4 | 10.47 |

